# No evidence for human monocyte-derived macrophage infection and antibody-mediated enhancement of SARS-CoV-2 infection

**DOI:** 10.1101/2020.12.22.423940

**Authors:** O García-Nicolás, P V’kovski, F Zettl, G Zimmer, V Thiel, A Summerfield

## Abstract

Vaccines are essential to control the spread of severe acute respiratory syndrome coronavirus-2 (SARS-CoV-2) and to protect the vulnerable population. However, one safety concern of vaccination is the possible development of antibody-dependent enhancement (ADE) of SARS-CoV-2 infection. The potential infection of Fc receptor bearing cells such as macrophages, would support continued virus replication and inflammatory responses, and thereby potentially worsen the clinical outcome of COVID-19. Here we demonstrate that SARS-CoV-2 and SARS-CoV-1 neither infect human monocyte-derived macrophages nor induce inflammatory cytokines in these cells, in sharp contrast to Middle East respiratory syndrome (MERS) coronavirus and the common cold human coronavirus 229E. Furthermore, serum from convalescent COVID-19 patients neither induced enhancement of SARS-CoV-2 infection nor innate immune response in human macrophages. These results support the view that ADE may not be involved in the immunopathological processes associated with COVID-19, however, more studies are necessary to understand the potential contribution of antibodies-virus complexes with other cells expressing FcR receptors.

## 1 Introduction

Since the emergence of the severe acute respiratory syndrome coronavirus-2 (SARS-CoV-2) in December 2019 in the Chinese city of Wuhan, the virus has spread globally causing the coronavirus disease 2019 (COVID-19) pandemic. The high morbidity and severity of COVID-19 in some of the affected patients have jeopardized the public health system of affected countries. In addition, the public health measures that have been implemented to control the pandemic have affected the life and economy of millions of people around the world.

In the current situation, there is no preexisting immunity to SARS-CoV-2, which allows the virus to spread rapidly in the human population. A global vaccination campaign may have the potential to finally control the pandemic and vaccination programs have started recently. However, a concern is the possibility that vaccination could promote antibody-dependent enhancement (ADE) of SARS-CoV-2 infection which could be associated with enhancement of the disease (Lee, Wheatley, Kent, & DeKosky, 2020)

The undergoing mechanism of ADE of infection is based on the interaction between virion-antibody complexes and Fc gamma receptors (FcγR) that are expressed by cells of the immune system such as macrophages. The binding of virion-antibody complexes to Fc receptors could result in their uptake into the cells by receptor-mediated endocytosis leading to potential infection of the cells (Taylor et al., 2015). Despite speculation and alarming about this possibility at the time of initiation of our work, no data were published specifically addressing ADE of SARS-CoV-2 (Lee et al., 2020; Rogers et al., 2020). With this in mind, the present study aimed to investigate whether immune sera from convalescent COVID-19 patients would enhance SARS-CoV-2 infection and promote secretion of pro-inflammatory cytokines production by human macrophages. To this end we performed a comparative study on the susceptibility of human macrophages to infection with human coronavirus 229E (HCoV-229E), Middle East respiratory syndrome coronavirus (MERS-CoV), SARS-CoV and SARS-CoV-2 and the inflammatory cytokine response of these cells. Potential ADE of infections by SARS-CoV and SARS-CoV-2 were studied using immune sera from convalescent COVID-19 patients.

## 2 Material and Methods

### 2.1 Ethics Statement

Buffy coats from anonymous healthy blood donors were obtained from the regional transfusion blood service of the Swiss Red Cross (SRC) (Bern, Switzerland). The use of buffy coats was approved by the SRC review board. All serum samples employed in this study were collected following the guidance of the Act on Medical Devices (MPG guideline 98/79/EC) for the collection of human residual material to evaluate suitability of an *in vitro* diagnostic medical device (§24). For this study an informed consent and ethical approval was not needed because only leftovers of serum samples for diagnostic laboratory procedures were used.

### 2.2 Cells

Vero cells (E6 and B4 lineages, African Green monkey kidney epithelial cells) and Huh-7 cells (human hepatocellular carcinoma cells) were cultured in Dulbecco’s minimal essential medium (DMEM; Life Technologies), supplemented with 10 % fetal bovine serum (FBS), non-essential amino acids (Life Technologies), penicillin-streptomycin (Gibco) and HEPES (Gibco).

For the production of human monocyte derived macrophages (hMDM), peripheral blood mononuclear cells (PBMCs) were isolated from buffy coats by density gradient centrifugation on Ficoll-Paque™ PLUS (1.077 g/L; GE healthcare). Then, monocytes were sorted using anti-CD14 beads as proposed by the manufacturer (Miltenyi Biotech GmbH), and seeded in 24 well plates at 2.5 × 10^5^ cells/well in 500 μl of Roswell Park Memorial Institute (RPMI) 1640 medium (Gibco) and kept at 37 °C and 5% CO_2_ atmosphere for one hour. Non-adherent cells were removed and 500 μl of RPMI 1640 supplemented with 10% of FBS (Gibco), GlutaMAX (Gibco), penicillin-streptomycin (Gibco) and human M-CSF (100 ng/ml; Miltenyi Biotec) were added. The cells were cultured for six days at 37 °C and 5% CO_2_. The full medium complemented with M-CSF was replaced every 48 to 72 hours.

### 2.3 Viruses

A collection of different coronaviruses was employed for the experiments of the present study including the human coronavirus 229E (HCoV-299E; (Thiel, Herold, Schelle, & Siddell, 2001)), Middle East respiratory syndrome coronavirus (MERS-CoV, strain EMC/2012; (van Boheemen et al., 2012)), SARS-CoV (Frankfurt-1; (Thiel et al., 2003)) and SARS-CoV-2 (SARS-CoV-2/München-1.1/2020/929) kindly provided by Daniela Niemeyer, Marcel Müller, and Christian Drosten (Charité, Berlin, Germany). HCoV-299E was propagated in Huh-7 cells in DMEM supplemented with 5% of FBS and non-essential amino acids at 33°C. MERS-CoV was propagated in Vero B4 cells in MEM supplemented with 2% of FBS and non-essential amino acids at 37°C. For the propagation of SARS-CoV and SARS-CoV-2 Vero E6 cells in MEM supplemented with 2% of FBS and non-essential amino acids at 37°C was employed. All viral titrations were performed taking advantage of the virus-induced cytopathic effects that were apparent 56 to 72 hours post infection. Virus titers were expressed as 50% tissue culture infective dose per ml (TCID_50_/ml).

### 2.4 Infection of hMDM

Human MDM were incubated for 1.5 h at 39°C or 37°C with the respective virus using a multiplicity of infection (MOI) of 1 TCID_50_ per cell, including mock control. Subsequently, the virus inoculum was removed, the cells washed three times with warm phosphate buffered saline (PBS), and RPMI medium supplemented with 2% FBS was added to the cells. As a positive control for the induction of pro-inflammatory cytokines either 1 μg/ml of lipopolysaccharide (LPS; Sigma-Aldrich) or 10 μg/ml of polyinosinic-polycytidytic acid (poly I:C, Sigma-Aldrich) were added to the cell culture medium. After 24 h, supernatants were collected and stored at −70°C. Cells were fixed with 4% formaldehyde for 10 min at room temperature,washed with PBS, and permeabilised with 0.3% saponin (PanReac AppliChem). The permeabilisation was performed for 20 min on ice in the presence of J2 monoclonal antibody direceted to dsRNA (English and Scientific Consulting). Subsequently, the cells were washed with 0.1% saponin, and cells were incubated for 20 min on ice with Alexa Fluor 488 conjugated anti-mouse IgG2a (ThermoFisher Scientific) in 0.3% saponin. Cells were washed once with PBS prior incubation for 5 min at 37 °C with 4′,6-diamidino-2-phenylindole (DAPI; Sigma). Finally, the percentage of infected hMDM was determined by by enumeratingof dsRNA positive cells in 10 fields/well using an Axio Observer Z1 inverted microscope equipped with a Zeiss Colibri Illuminator (CarlZeiss) and digital imaging Zeiss software (AxioVision, v4). All generated images were analyzed using ImageJ software.

### 2.5 Antibody-Dependent Enhancement of Infection

A collection of sera from COVID-19 convalescent patients from a previously published work was employed for the present study (Zettl et al., 2020). This included sera with a broad range of neutralization titers against SARS-CoV-2 (ND_50_ <1:10; 1:20; 1:160; 1:240 and 1:2560). In order to test the ADE potential of these sera, different serum dilutions (1:10; 1:100; 1:1000 and 1:10000) were incubated for 30 min at 37 °C with an equal volume of viral suspension (SARS-CoV or SARS-CoV-2) corresponding to a MOI of 1 TCID_50_/cell. Thereafter, the virus/serum mixtures were added to human macrophages or Vero E6 cells and incubated for 30 min at 37 °C and 5% CO_2_ atmosphere. The cells were washed three times with PBS before fresh medium was added. Following an incubation period of 24 h, macrophages and Vero E6 cells were analyzed by flow cytometry for SARS-CoV-2 and SARS-CoV nucleocapsid (N) protein expression. The cells were harvested using TrypLE™Select (Gibco) for 20 min at room temperature and fixed with 4% (w/v) formaldehyde. Thereafter the cells were washed and permeabilized/stained for 20 min on ice with 0.3% (*w*/*v*) saponin in PBS in the presence of a rabbit antibody directed to the SARS-CoV N protein (Rockland-inc). The cells were subsequently washed, incubated for 10 min with anti-rabbit Alexa 488 fluochrome conjugate (ThermoFisher Scientific), and analysed by flow cytometry using a FACSCantoII (Becton Dickinson). For analysis, Flowjo V.9.1 software (Treestars, Inc.) was used. Dead cells were excluded by electronic gating in forward/side scatter plots, followed by exclusion of doublets.

### 2.6 Cytokines Measurement

Cytokines in the supernatants in this study including tumor necrosis factor (TNF), interferon beta (IFN-β), interleukin 6 (IL-6) and IL-1β were quantified by ELISA (R&D Systems) following the manufacturer’s instructions. Detection limits were 30 pg/ml for TNF, 10 pg/ml for IFN-β, 4 pg/ml for IL-1β and 9 pg/ml for IL-6.

### 2.7 Statistics

For the generation of figures and data analyses the GraphPad Prism 8 Software (GraphPad Software, Inc.) was employed. All experiments were independently performed 3 to 6 times with cells from different donors, and each experiment was run in triplicates. For viral titrations, differences between groups were assessed using a Kruskal–Wallis test, and for individual differences the Mann–Whitney *U*-test with Bonferroni correction as *post hoc* was employed. For group differences in the percentages of infected cells and levels of cytokines expression comparisons, a one-way ANOVA test with Bonferroni correction as *post hoc* was performed. Correlation analysis between infected cells, viral titers, and expressed cytokines were calculated by Spearman's Rho analysis; a correlation between two different factors was considered relevant with R^2^ > 0.5. A *p* value lower than 0.05 was considered statistically significant for every analyzed data. In figure 1, different superscript letters indicate a significant difference (p < 0.05) between the conditions. For the table 2 ^*^ indicates *p* < 0.05, ^**^*p* ≤ 0.002, ^***^*p* ≤ 0.001 and ^****^*p* ≤ 0.0001.

**Figure 1.**
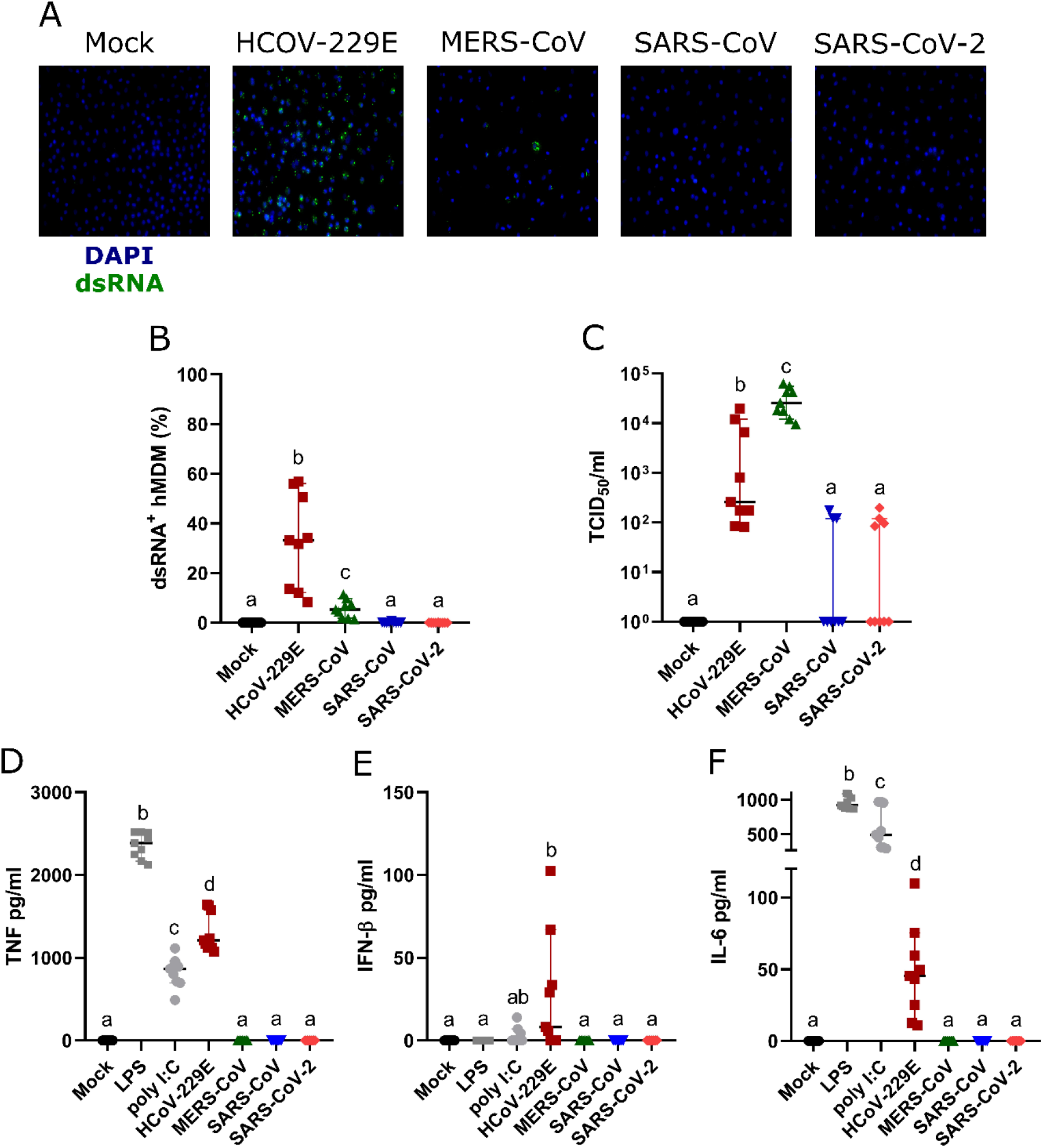
Susceptibility of hMDM to different human coronaviruses. Human MDM were inoculated with different coronaviruses (hCoV-229E, MERS-CoV, SARS-CoV-1 and SARS-CoV-2) using an MOI of 1 TCID_50_/cell. Mock-infected cells or cells treated with LPS or poly I:C served as controls. After incubating the cells for 1.5 hours, the inoculum was removed, the cells washed, and fresh medium added. At 24 hours post infection, dsRNA in the cells was detected with a specific antibody and nuclei were stained with DAPI (A). The percentage of dsRNA-positive hMDM was calculated for 10 fields per condition (B). Virus titers (C), TNF-α (D), IFN-β (E) and IL-6 (F) were determined in the cell culture supernatants. The data from three independent experiments run in triplicates are shown. Different superscript letters indicate a significant difference (p < 0.05) between the conditions.

## 3 Results

### 3.1 Human coronaviruses differ in their ability to infect hMDM

Infection of hMDM with HCoV-299E, MERS-CoV, SARS-CoV and SARS-CoV-2 at MOI of 1 TCID_50_/cell demonstrated high susceptibility to infection with the common cold virus HCoV-229E, low susceptibility to infection with the highly pathogenic coronavirus MERS-CoV, and resistance to infection by SARS-CoV and SARS-CoV-2, all assessed by immunofluorescence analysis using an anti-dsRNA marker (Figure 1A and B). When we quantified the number of infectious virus particles in the cell culture supernatants we observed that only HCoV-229E and MERS-CoV replicated efficiently in hMDM (Figure 1C). Although HCoV-229E showed higher percentages of infected hMDM (32.79% ±18.79 SD) the highest virus titers were found in the cell culture supernatant of macrophages infected with MERS-CoV (Figure 1C). Nevertheless, it has to be taken into consideration that all experiments were performed at 37°C although the optimal temperature for HCoV-229E is 33°C (Dijkman et al., 2013). Viral titers of SARS-CoV and SARS-CoV-2 were not statistically significant compared to the mock control. In conclusion, these results indicate that neither SARS-CoV nor SARS-CoV-2 are able to infect hMDM.

### 3.2 Human MDM only produce cytokines after HCoV-229E infection

Human MDM infected by HCoV-229E, but not MERS-CoV, SARS-CoV and SARS-CoV-2, secreted TNF, IFN-β and low levels of IL-6 (Figure 1D-F). None of the tested coronaviruses induced secretion of IL-1β. Taking into consideration that viral RNA might induce innate immune responses, we tested the correlation between the percentage of dsRNA positive cells, viral titers and level of secreted cytokines. The results found a clear association between the percentage of infected cells and the secreted cytokines but not with the viral titers (Table 1).

**Table 1.**
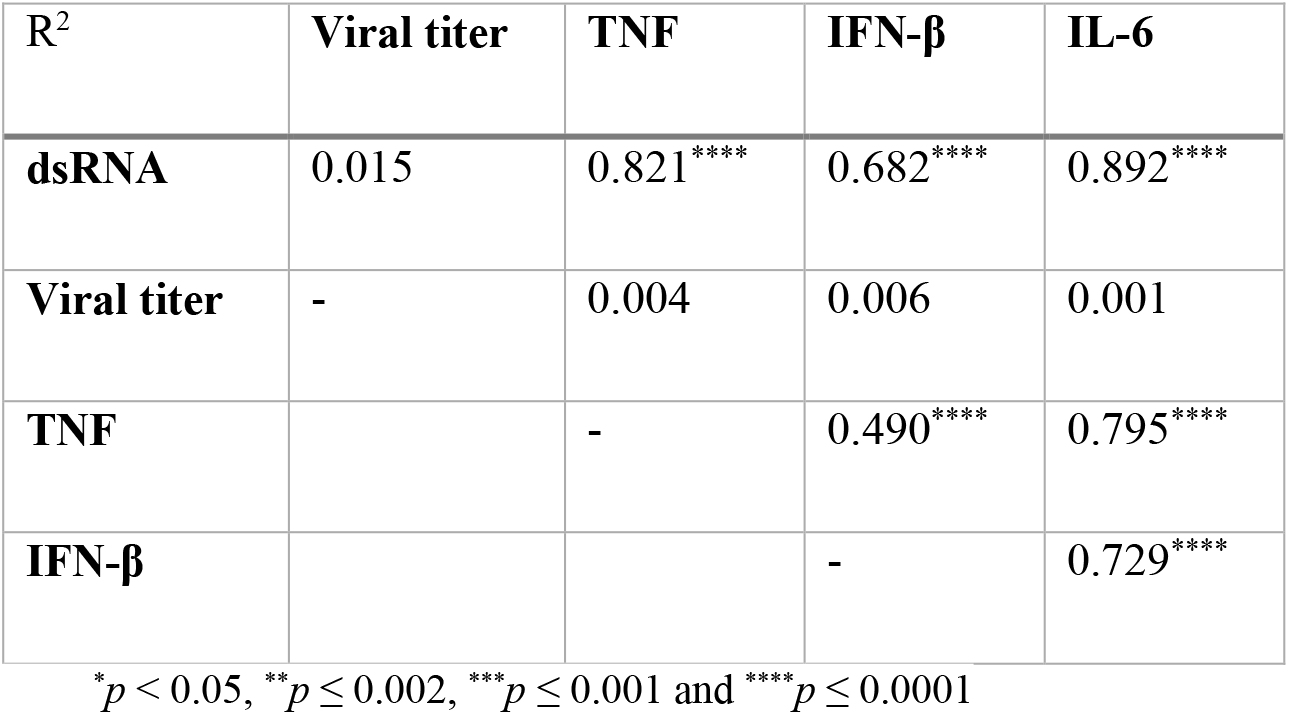
Correlation analysis between dsRNA positive hMDM, infectious virus titers and pro-inflammatory cytokines

| R<sup>2</sup> | Viral titer | TNF | IFN-β | IL-6 |
| --- | --- | --- | --- | --- |
| **dsRNA** | 0.015 | 0.821**** | 0.682**** | 0.892**** |
| **Viral titer** | - | 0.004 | 0.006 | 0.001 |
| **TNF** |  | - | 0.490**** | 0.795**** |
| **IFN-β** |  |  | - | 0.729**** |

\* $p < 0.05$ , \*\* $p \leq 0.002$ , \*\*\* $p \leq 0.001$ and \*\*\*\* $p \leq 0.0001$

### 3.3 Antibodies from convalescent COVID-19 patients neither induce antibody-dependent enhancement of infection of hMDM with SARS-COV-2 nor promote cytokine responses

In order to control the experimental conditions for testing ADE, selected COVID-19 sera with a neutralization titer below 1:10 and with a neutralization titer of 1:240, were diluted serially from 1:10 to 1:1000, mixed with SARS-CoV and SARS-CoV-2, and then added to Vero E6 cells. Both sera demonstrated a dilution-dependent inhibition of SARS-CoV-2 infection. In addition, a cross-reactivity of COVID-19 convalescent patient sera with SARS-CoV-1 was observed, confirming previous reports (Zettl et al., 2020) (Table 2).

Using the same approach, a larger collection of sera from COVID-19 convalescent patients with neutralization titers ranging from <10 to 1:2560 was incubated at different concentrations with SARS-CoV and SARS-CoV-2 and added to hMDM. Importantly, with none of the tested serum dilutions which went up to1:10’000 infection of hMDM was observed. Moreover, hMDM exposed to virus-antibodies complexes did not secrete any detectable pro-inflammatory cytokines (Table 2). These results indicate that the potential uptake of SARS-CoV-1 and SARS-CoV-2 via FcR does not result in infection and activation of human macrophages.

**Table 2.**
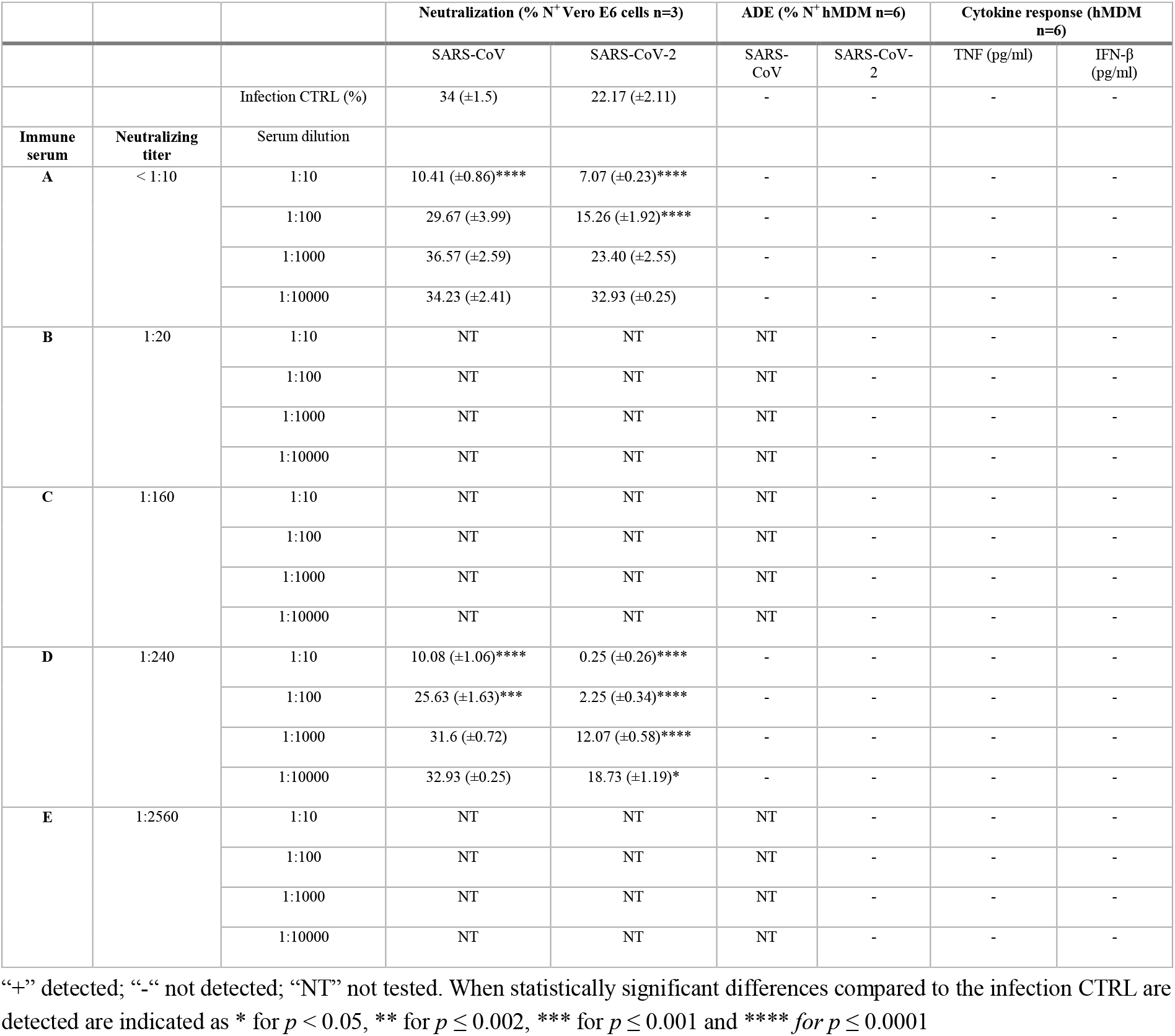
Summary data for neutralization on Vero E6 cells and antibody dependent enhancement of infection in or cytokine response in hMDM for SARS-CoV and SARS-CoV-2-antibody complexes

|  |  |  | Neutralization (% N<sup>+</sup> Vero E6 cells n=3) |  | ADE (% N<sup>+</sup> hMDM n=6) |  | Cytokine response (hMDM n=6) |  |
| --- | --- | --- | --- | --- | --- | --- | --- | --- |
| | | | SARS-CoV | SARS-CoV-2 | SARS-CoV | SARS-CoV-2 | TNF (pg/ml) | IFN-β (pg/ml) |
| | | Infection CTRL (%) | 34 (±1.5) | 22.17 (±2.11) | - | - | - | - |
| Immune serum | Neutralizing titer | Serum dilution |  |  |  |  |  |  |
| **A** | < 1:10 | 1:10 | 10.41 (±0.86)**** | 7.07 (±0.23)**** | - | - | - | - |
| | | 1:100 | 29.67 (±3.99) | 15.26 (±1.92)**** | - | - | - | - |
| | | 1:1000 | 36.57 (±2.59) | 23.40 (±2.55) | - | - | - | - |
| | | 1:10000 | 34.23 (±2.41) | 32.93 (±0.25) | - | - | - | - |
| **B** | 1:20 | 1:10 | NT | NT | NT | - | - | - |
|  |  | 1:100 | NT | NT | NT | - | - | - |
|  |  | 1:1000 | NT | NT | NT | - | - | - |
|  |  | 1:10000 | NT | NT | NT | - | - | - |
| **C** | 1:160 | 1:10 | NT | NT | NT | - | - | - |
|  |  | 1:100 | NT | NT | NT | - | - | - |
|  |  | 1:1000 | NT | NT | NT | - | - | - |
|  |  | 1:10000 | NT | NT | NT | - | - | - |
| **D** | 1:240 | 1:10 | 10.08 (±1.06)**** | 0.25 (±0.26)**** | - | - | - | - |
| | | 1:100 | 25.63 (±1.63)*** | 2.25 (±0.34)**** | - | - | - | - |
| | | 1:1000 | 31.6 (±0.72) | 12.07 (±0.58)**** | - | - | - | - |
| | | 1:10000 | 32.93 (±0.25) | 18.73 (±1.19)* | - | - | - | - |
| **E** | 1:2560 | 1:10 | NT | NT | NT | - | - | - |
|  |  | 1:100 | NT | NT | NT | - | - | - |
|  |  | 1:1000 | NT | NT | NT | - | - | - |
|  |  | 1:10000 | NT | NT | NT | - | - | - |

## 4 Discussion

A first observation of the present study was that in contrast to the common cold virus hCoV-229E and MERS-CoV, SARS-CoV and SARS-CoV-2 were unable to infect hMDM. The receptor for SARS-CoV-2 is angiotensin-converting enzyme 2 (ACE2) typically expressed on ciliated epithelial cells, goblet cells, type II alveolar pneumocytes as well as other cells from different organs as enterocytes (Sungnak et al., 2020). However, there are conflicting reports on the infection of human macrophages by SARS-CoV. While one study described very limited ACE2 expression by macrophages (Sungnak et al., 2020), another report postulated that the receptor is expressed on tissue resident macrophages (Song et al., 2020). In COVID-19 patients, the viral nucleoprotein N has been detected in macrophages from lymphoid organs of COVID-19 patients (Park, 2020), but it is not clear whether this was caused by direct infection or as a consequence of phagocytosis of infected cells.

The infection of hMDM by HCoV-229E is in line with the expression by these cells of aminopeptidase N (CD13), the cellular receptor for HCoV-229E (Yeager et al., 1992). It is also in agreement with previous reports describing the infection of alveolar macrophages by HCoV-229E (Funk et al., 2012). The cellular receptor for MERS-CoV dipeptidyl peptidase-IV (DPP4 also known as CD26) is expressed at low levels by human monocytes and macrophages of healthy donors (Rao et al., 2018; Rao et al., 2019; Wang et al., 2013; Zhong et al., 2013), which could explain MERS-CoV infection of hMDM in our experiments.

Our results are also in line with a previous report demonstrating that following infection with HCoV-229E human macrophages strongly secrete TNF, but also produce IL-6 and some IFN-β (Funk et al., 2012). Finally, we also noticed a poor innate immune response of macrophages following infection with MERS-CoV, confirming a previous report (Zhou et al., 2014).

While writing the present manuscript contradictory data were published claiming that SARS-CoV-2 induces an immune activation of hMDM (Zheng et al., 2020). These conflicting results might be the consequence of different methodologies used. While we employed purified monocytes to generate macrophages in six days, Zheng and collaborators kept PBMC with M-CSF for four days and then for another 10 days when the cells became adherent (Zheng et al., 2020). Another important methodological difference is that our study used ELISA to detect cytokines while Zheng and collaborators analyzed mRNA by RT-PCR. As the levels of mRNA induction found were rather low (fold change increase below 0.3), it is possible that protein detection by ELISA would have been below the detection limit as well. We are therefore proposing that hMDM generated from pure monocytes are not permissive to SARS-CoV-2 infection and do not mount inflammatory responses. This is in contrast to HCoV-229E and the TLR ligand controls. While future studies expanding to tissue macrophages are important, our results indicate that proinflammatory responses observed during COVID-19 may not be the result of macrophage infection but rather originate from other innate immune cells or a complex interaction between different immune cells which could include macrophages. Therefore, to understand these events more immune cells such as plasmacytoid dendritic cells that are at the frontline of the antiviral cytokine responses should be investigated.

In view of a potential link between ADE and inflammation during COVID-19 in the presence of antibodies, we tested this hypothesis using hMDM. With the selected sera from convalescent COVID-19 patients and the described conditions, even at very high serum dilutions and with sera that had low levels of neutralizing antibodies, we did not find evidence for antibody mediated enhancement of macrophage infection or pro-inflammatory cytokine responses. Interestingly, sera of COVID-19 convalescent patients showed cross-reactivity to SARS-CoV (Zettl et al., 2020) but did not enhance infection by this virus of macrophages SARS-CoV, although having cross-reactivity. This is in line with a previous report showing that the presence of cross-reacting antibodies against SARS-CoV-2, originating from previous endemic coronavirus infections, were not linked with more severe COVID-19 (Ng et al., 2020). Furthermore, in vaccination/challenge experiments carried out in macaques no signs of enhanced disease were detected (Gao et al., 2020; Yu et al., 2020). Finally, COVID-19 patients treated with plasma transfusion from convalescent patients did not show signs of disease aggravation (Casadevall & Pirofski, 2020; Joyner et al., 2020). Altogether, these data are in line with our findings, demonstrating that hMDM are not infected or activated by SARS-CoV-2 neither by direct contact nor mediated by antibodies from convalescent COVID-19 patients. Although the lack of hMDM infection by SARS-CoV and SARS-CoV-2 provides evidence of a lack of ADE, our study cannot exclude ADE effects on other FcR-expressing cells as well as a possible role of the complement system or T-cell mediated inflammation.

## Acknowledgments

We would like to thank all members of the IVI and for their support and helpful discussions. We are grateful Daniela Niemeyer, Marcel Müller, and Christian Drosten (Charité, Berlin, Germany) for providing the viruses and to the Swiss Transfusion SRC (Swiss Red Cross) Inc. (Regional transfusion blood service, Bern, Switzerland) for providing human buffy coats.

